# Early lethality of embryos derived from transgenic *Xenopus* females is associated with reduced ovarian *grem1* expression

**DOI:** 10.1101/268235

**Authors:** Caroline W. Beck, Joanna Ward, Lisa Troise, Catherine Brochard

## Abstract

**The *grem1* gene codes a protein that inhibits the action of multiple members of a growth factor family known as bone morphogenetic proteins (BMPs). Certain members of this BMP family can regulate both fecundity and fertility in mammals via their action on oocyte (egg) development, and *grem1* has been identified as a marker of oocyte quality in humans. The model amphibian *Xenopus laevis* is far more fecund than mammals, producing thousands of eggs in a clutch. However, female transgenic frogs carrying *grem1* under the control of a stress inducible *hsp70* promoter (“G” frogs) produce very few viable offspring. Here, we show that this is not due to reduced fecundity or fertilization rate, but results from a significant reduction in subsequent survival over the first day of development. Embryos that successfully survive for the first day were found to go on to develop normally when compared to their peers. Both the morphology and stage distribution of oocytes from G females appears normal, and oocytes develop at expected rates, although stage VI oocytes were found to have a lower response to in vitro progesterone treatment. Unexpectedly, levels of *grem1* mRNA were found to be consistently lower in the female ovaries from four independent G transgenic lines than in wild type ovaries. Both transgenic and wild type offspring were equally affected, confirming a maternal effect. Our study shows that transgenic females with the lowest levels of *grem1* transcripts in the ovary have the lowest rates of survival past the first day of amphibian embryogenesis, equivalent to pre-implantation staged mammalian embryos. The reduced expression of *grem1* in the oocytes of transgenic females suggests transgene supression of an endogenous locus may occur in the *Xenopus* female germline, an unexpected finding.**

## Introduction

Infertility, which affects 70 million human couples worldwide, can have a genetic basis (1). Female infertility can result from defective development of the oocyte, and two of the very few genes associated with this kind of infertility belong to the same family of growth factors: the bone morphogenetic proteins (BMPs). In humans, both BMP15 (2, 3) and its close relative GDF9 (growth and differentiation factor 9) (4) are implicated in fertility. Both GDF9 and BMP15 are actively transcribed in developing oocytes, although not exclusively. In ewes, heterozygous mutations in GDF9 or BMP15 can increase fecundity by increasing the rate of ovulation, but homozygous mutations result in infertility (5-7).

BMP15 and GDF9 are therefore targets for fertility regulation in sheep and cattle (8). However, in mice, which have larger litters, while GDF9 loss results in infertility (9) loss of BMP15 only reduces litter size (10). As well as GDF9 and BMP15 in the oocytes, BMP4 and BMP7 are expressed in the surrounding stroma during oogenesis and positively regulate oocyte development. BMPs are secreted proteins that bind as dimers to type I and II receptor serine/threonine kinases in the plasma membrane of target cells and elicit cell signalling via activation of a group of cytoplasmic proteins known as SMADs (11). Together these form transcriptional complexes, which regulate specific gene expression at BMP response elements (12).

*Xenopus laevis*, the south African clawed frog, is a useful model for the study of development as eggs can be obtained all year round following injection with human chorionic gonadotrophin (HCG), and these are large enough to be easily manipulated, with a diameter of 1.2 mm. Oogenesis, the progression from primordial germ cell to mature egg, is an asynchronous process in *Xenopus*, with all stages being present in the ovary throughout adult life. Oogenesis results in the formation of a polarised egg in *Xenopus* and six stages of oocyte development (I-VI) were identified and described by Dumont (13). The germ cells (oogonia) first undergo a series of mitotic cell divisions leading to the formation of the primary oocyte. Meiosis then begins, but is arrested in prophase I. The oocyte then undergoes a period of growth, and uptake of nutrients, such as vitellogenin: Dumont stages II to V are considered vitellogenic, stage VI is post-vitellogenic. Animal-vegetal polarity is established as pigment is displaced to one pole and mitochondria to the other. In the response to HCG, progesterone is produced from ovarian follicle cells and induces the resumption of meiosis in stage VI oocytes. The germinal vesicle (nucleus) migrates to the animal pole where it is visible as a white spot, forming a visible marker of the process, and the first polar body is lost. The resulting mature oocyte (egg) is arrested at metaphase II pending fertilization. Once this occurs, the egg completes meiosis and the second polar body is lost.

BMPs are potent morphogens, able to elicit distinct cellular responses at different concentrations, and so their dispersal is regulated by a group of extracellular inhibitors (11). *grem1* encodes Gremlin1, a cysteine knot protein that acts as an extracellular inhibitor of several members of the bone morphogenetic protein (BMP) family (14). Gremlin1 binds directly to BMPs preventing them activating their receptors. In our laboratory, we have generated several lines of transgenic *Xenopus laevis* in order to study the role of Gremlin1 in limb development and regeneration (15). These animals carry the double transgene (Fig 1a). Animals carrying this transgene are known as G, and each founder has a unique number (Fig 1b,c). These animals can be “genotyped” by the presence of red fluorescent protein (RFP) in the lens of the eye. While they do not express *grem1* from the transgene heat shock protein 70 (hsp70) promoter at ambient temperatures, following a heat shock (transfer of animals to 34°Cfor 30 minutes), they rapidly express *grem1* in all cells (15). However, even at ambient temperatures, female frogs carrying the G transgene were unexpectedly observed to exhibit reduced fertility, so these previous studies relied on the use of male carriers. Intriguingly, *grem1* had already been associated with fertility in humans. Increased *grem1* transcription in human cumulus granulosa cells (GC) is associated with better blastocyst formation, embryo quality and improved pregnancy success rate (16, 17). The aim of the study was to findthe underlying cause of reduced viable offspring from G mothers and to determine how the transgene insertion could cause this.

## Methods

### Ethics approval

All protocols involving animals used in this study were approved by the University of Otago’s Animal ethics committee under protocol AEC97/15.

### Transgenic animals and crosses

G transgenic founder frogs contain a random insertion of the *hsp70:grem1*,γ-*crystallin:RFP* (G) transgene (Fig 1a). Transgenic animals used in this study were generated using sperm nuclear injection for a previous project, and their design is described therein (15). Adult females were between 2 and 6 years old at the time of the study. Adult males were between 1 and 6 years old. As founders are produced by titrated random transgene insertion into fertilised eggs, these animals are expected to be heterozygous for the transgene. Five male founders (G5, G6, G7, G8 and G14) were crossed to a single wild-type (WT) female as well as to individual WT females to determine paternal effects (Fig 1b). Three female founders (G3, G9 and G12) and four F1 females (G8.x) descended from male founder G8 were used to determine maternal effect and transgene position effect, by crossing to the same WT male (Fig 1c).

### Fecundity assay

Adult female *Xenopus laevis* were first weighed and then induced to ovulate by injecting 6.7 U HCG/g bodyweight (i.e. 500U for a 75g female) into the dorsal lymph sac. Injected females were housed in pairs in an incubator at 17°Cfor 16 hours overnight. In the morning, females were individually transferred to 1x MMR (Marc’s modified ringers). Once egg laying had commenced, the number of eggs laid in one hour was counted for each individual. Due to the strong positive correlation between WT female bodyweight before induction and the number of eggs laid in one hour (Fig 2a), the data were normalised for each animal by dividing raw counts by bodyweight in grams and multiplying by 62g, the median frog weight in this study.

**Fig. 1.**
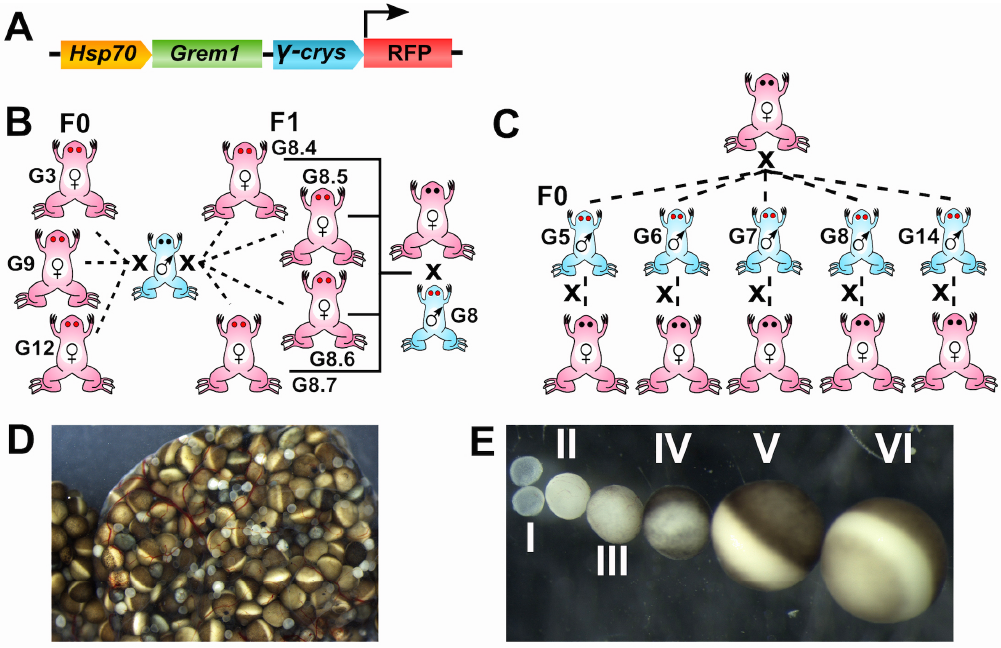
Design of transgene and parent of origin crossing scheme and oocyte staging. A) Schematic of G transgene as published and characterised in (15). At the temperatures used to house frogs and raise embryos, the *hsp70* promotor is expected to be silent (15), so no transgene derived *grem1* transcripts are detectable. The γ-crystallin promotor drives expression of red fluorescent protein in the lenses of G tadpoles and frogs, allowing easy genotyping. B) Back crossing scheme used for transgenic female frogs. Founders G3, G9 and G12 as well as 4 female descendants of male founder G8 were crossed to the same wild type (WT) male frog. C) Back crossing scheme used for transgenic male frogs. Founders G5, G6, G7, G8 and G14 were crossed both to the same WT female and also to individual WT females. All founders in this study were heterozygous for the G transgene, but have random insertion sites. D) an ovarian lobe from an uninduced female wild type *Xenopus*. Note all stages are present at once. E) Collagenase treated oocytes staged from I to VI as in (13).

### *In vitro* fertilisation rate

All *in vitro* fertilisation was done using naturally laid eggs, with individual females housed in 1x MMR as above, to prevent activation. Sperm suspension was prepared by gently homogenising the testes from a euthanized male frog in 1ml 1 × MMR using a plastic pestle. Approximately 100 eggs were transferred to 50mm petri dishes with minimal 1 × MMR and fertilised by adding 20l sperm suspension, waiting 1-2 minutes, and then flooding eggs with water. Three replicates of each fertilisation were performed. Eggs were scored as fertilised once the embryo had begun to cleave, and fertility was defined as the percentage of eggs reaching the 2-4 cell stage.

### Determination of rate of survival to 24 hours

Following fertilisation confirmation, embryos were treated with 2% Cysteine HCl pH 7.9 in 0.1 × MMR to completely remove the jelly coats, rinsed 3 × with 0.1 × MMR, and the unfertilised eggs disregarded. Remaining embryos were incubated at 18°Covernight. 28 hours after initial fertilisation, which equates to completion of gastrulation/start of neurulation at this temperature, or Nieuwkoop and Faber stage 14 (18), the number of embryos developing normally in each dish were counted. Counts were also obtained for embryos that had died or undergone abnormal gastrulation, before discarding these. Percentages of normal, abnormal and dead embryos were calculated relative to the number of successfully fertilised eggs, with three replicates for each cross.

### Determination of transgenic ratios and survival to six days

The number of transgenic vs. non-transgenic survivors was determined at day 6 after fertilisation, by observing presence or absence of the reporter red fluorescent protein (RFP) in the lenses of the embryonic eyes. As all crosses were heterozygote x wild type (i.e. back-cross) regardless of the sex of the transgenic animal, half the offspring are expected to carry the transgene. Chi squared tests were used to determine any variation from the expected Mendelian ratio of RFP to non-RFP animals (50:50). Percentages of surviving, transgenic and non-transgenic embryos were calculated relative to the number of successfully fertilised eggs with three replicates for each cross.

### Long-term survival analysis

Embryos developing normally at 6 days post fertilisation were first sorted by transgenic status and the three replicate samples pooled before transferring to tanks to a recirculating aquarium. Daily feeding with spirulina and ground salmon pellets commenced at this stage. The aquarium was kept at 23°C. Development to 10 weeks post fertilisation was assessed after terminal anaesthesia (MS222 overdose) by recording the developmental stage of each individual tadpole using the staging system of Nieuwkoop and Faber (18).

### Ovary composition and oocyte counting

Female *Xenopus* were sacrificed and their ovaries examined after 6 weeks of recovery following HCG induction. A single lobe from each ovary was isolated for each female and placed in a sealed 15 ml tube with 10 ml 0.2% collagenase in 1x MMR to separate the oocytes. Collagenase treatment was performed in a 30°Cincubator with constant slow end-over-end rotation. After 30 minutes, oocytes could be separated from each other and counted by stage according to Dumont (13).

### Oocyte maturation assay

For each female in the study, a total of 20 Stage VI oocytes that had been isolated from collagenase treated ovaries (10 from the left and 10 from the right) were exposed to 1 × MMR containing either 30μM progesterone (prepared as 30mM stock in ethanol and used fresh) or vehicle for 10 minutes. The oocytes were then gently rinsed in 1 × MMR and transferred to agar lined petri dishes containing 1 × MMR containing penicillin/streptomycin for 4 hours to overnight. The breakdown of the germinal vesicle (female nucleus) leads to a visible white spot appearing at the animal pole and indicates successful maturation of the oocyte. Oocytes were scored as mature, immature or dead.

### Quantitative real-time PCR (qRT-PCR.)

Whole ovary tissue was homogenised in TRIzol and total RNA isolated. RNA was subjected to DNase treatment and cDNA was generated using MMLV reverse transcriptase and oligo dT18. Gene-specific primers for qPCR were designed using NCBI primer designer to *X. laevis* transcript reference sequences or as described on Xenbase (Table 1). Tm of primers was between 63.6 - 69.5°C. Where possible, primers validated previously were used. A serial dilution of cDNA was used to generate a standard curve for each primer pair to determine primer efficiency. Sample cDNA was diluted to fall within the range of this standard curve. qPCR was performed on a Quantstudio 5 (Applied Biosystems) platform using 2 × SYBR green master mix (Takara Premix Ex Taq). Briefly, a mastermix was made, containing 0.5 μM of each of the corresponding forward and reverse primers and 5 μl of 2 × SYBR green mastermix (Takara). 9l of this mastermix was added to 1l of diluted cDNA in a 96-well qPCR plate. Cycling conditions were: 95°C2 min followed by 40 cycles at 95°C, 5 s, 62°C, 10 s, 72°C5 s. Reactions were set up in duplicate for each cDNA and target gene and all samples were normalised to the geometric mean of *Gapdh* and *Ef1-α* within the same sample to correct for efficiency of reverse transcription. Note that, for oocytes, only *Gapdh* was used for normalisation. The expression levels of genes of interest were calculated using Quantstudio Design and Analysis software (Applied Biosystems), using the standard curve method to normalise the amplification efficiency of target and reference genes as well as run-to-run differences. Specificity of the qPCR was confirmed by the melting curve of amplified products. Each qPCR experiment included a standard curve of each primer set to determine primer efficiency. Primer efficiency lay between 90.7 - 109%. This efficiency was used in the normalisation and fold change calculation.

### Data analysis

All graphs were produced and statistical analyses performed as described using Graph Pad Prism 7.0 software.

## Results

### The *grem1*transgene G does not significantly alter fecundity of female *Xenopus*

The number of eggs laid in one hour was calculated for each female in the study. Due to the strong correlation between female bodyweight before induction and the number of eggs laid in 1 hour (Table 2, Fig 2a), the data were normalised for pre-induction body weight. While G female founders typically laid fewer eggs per hour than either G8 F1 or wild type females (Fig 2b), a one-way ANOVA test showed no significant differences between means (F=0.69; p=0.524) of founder G females, G8 F1 females, and wild type animals when egg laying rate was normalised for bodyweight.

### Neither maternal nor paternal *grem1* transgene G alters fertility of *Xenopus* eggs

We sacrificed five transgenic founder males confirmed to be carrying the G transgene and carried out in vitro fertilisation of eggs from a single female WT frog, as well as to different WT females (Fig 1b). The rate of fertilisation was calculated as a percentage of eggs reaching 2 to 4 cell stage, for three replicate fertilisations of around 100 eggs each. For male founders back-crossed to the same individual WT female, fertility ranged from 66% (G6 male) to 93% (G8 male), with a wild type control male giving 87% with the same female. A one-way ANOVA showed no significant difference in mean fertilisation rate amongst males (F= 2.53; p=0.09; Fig 2c). Similarly, when eggs from female founders (n=3), female F1 derived from G8, and wild type females were fertilised with sperm from the same WT male (Fig 1c), one way ANOVA again showed no significant difference in mean fertilisation rate, (F=2.58, p=0.07, Fig 2d).

**Table 1.**
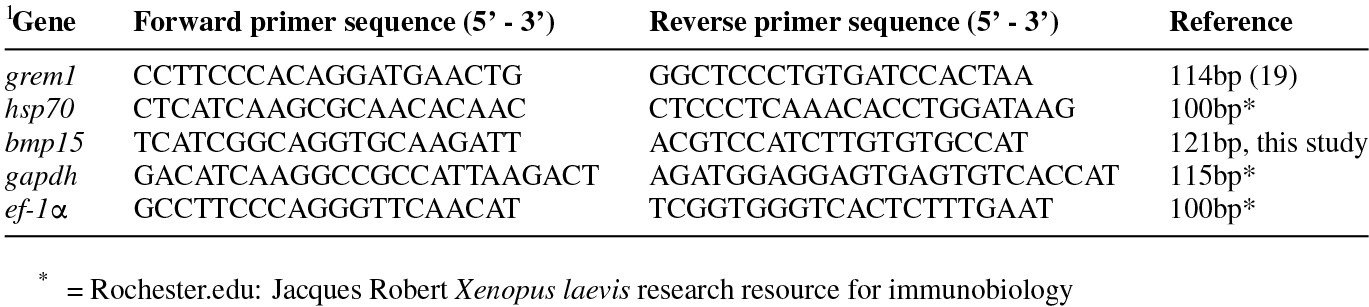
Primer Sequences

**Fig. 2.**
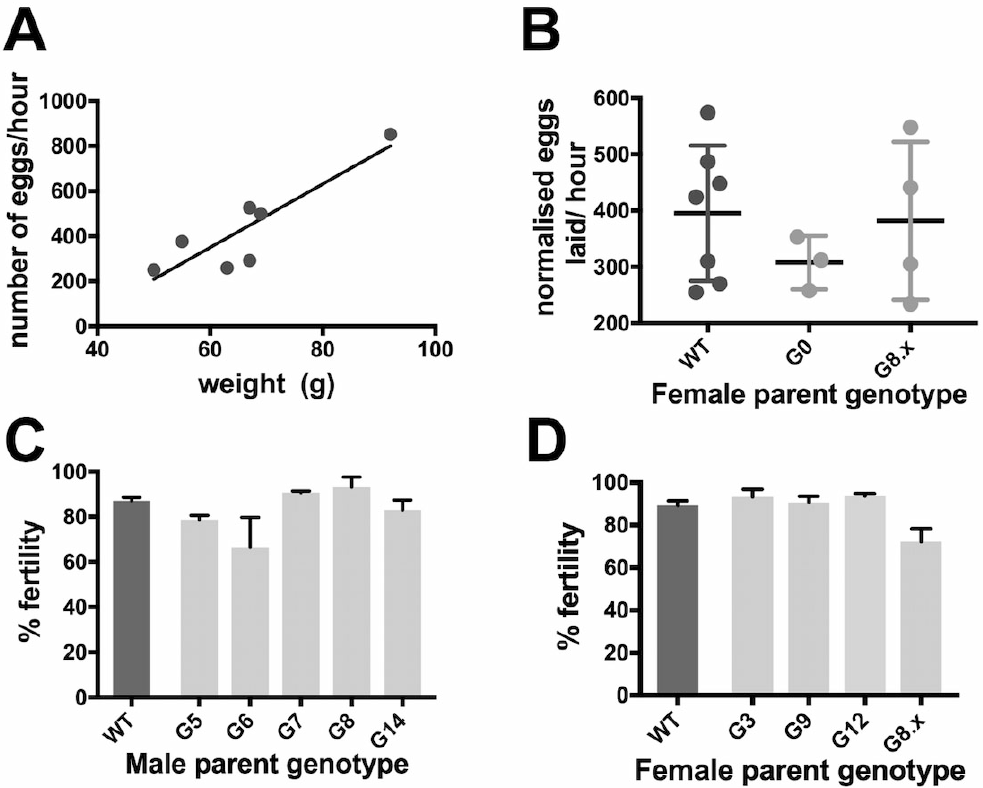
There is no effect of parental transgenic genotype on fecundity or fertilisation rate. A) linear regression analysis of seven WT females in the study showing positive correlation between egg laying rate and bodyweight on induction. B) Scatter plot comparing fecundity of three female G founders (G3, G9, G12), four female G8 F1 and seven wild type female *Xenopus*. Centre lines show the mean egg laying rate (per hour, normalised for body weight), and error bars are SD. No significant differences were seen between the three groups. C) Column graph showing that mean percentage successful fertilization rate was not altered when the male parent carried the G transgene. Sperm from five different founder G males and one WT male was used to fertilize 3 batches of around 100 eggs from the same WT female and fertilization scored as the percentage of eggs undergoing successful progression to 4 cells. Error bars are SEM. One-way ANOVA revealed no differences in fertility between any of the male parent genotypes. D) Column graph showing mean percentage fertilization was not affected when the female parent carried the G transgene. Sperm from the same WT male was used to fertilise 3 batches of around 100 eggs from each of two WT, three G founders and 4 G F1 females derived from the G8 male. One-way ANOVA revealed no differences in fertility between any of the female parent genotypes.

**Table 2.** Rate of egg laying in G founder, G8 F1 and wild type females

| Female | Pre-induction weight (g) | Eggs laid /hour | Eggs/hour normalised for bodyweight | normalised mean (+/-SD) |
| --- | --- | --- | --- | --- |
| G3 | 67 | 381 | 353 | 307 (+/- 48) |
| G9 | 39 | 162 | 258 |  |
| G12 | 62 | 312 | 312 |  |
| G8.4 | 49 | 433 | 433 | 382 (+/- 140) |
| G8.5 | 62 | 305 | 305 |  |
| G8.6 | 51 | 363 | 441 |  |
| G8.7 | 77 | 291 | 234 |  |
| WT1 | 50 | 250 | 310 | 395 (+/- 120) |
| WT2 | 67 | 292 | 270 |  |
| WT3 | 63 | 259 | 255 |  |
| WT4 | 92 | 852 | 574 |  |
| WT5 | 67 | 526 | 487 |  |
| WT6 | 55 | 376 | 424 |  |
| WT7 | 69 | 499 | 448 |  |

### Embryos with a *grem1* mother are less likely to survive earlyembryonic development

Having found no differences in fecundity or fertility to explain the difficulty breeding G female frogs, we next examined early development. *Xenopus* embryos are dependent on maternal transcripts until the mid-blastula transition at stage 8, so we would expect any maternal effect to impact early on in the animal’s development. We analysed the percentage of embryos that had been successfully fertilised from each replicate as either normal, dead or poorly gastrulated at stage 14. Survival rates (normal developing %age) of embryos derived from G females ranges from 3% to 59%, with wild types at 73 to 92%. Generally, mean survivability was lowest in offspring of females carrying G, but there were also differences depending on the founder, indicating that transgene position and context may be important.

One-way ANOVA showed significant variation (F=5.35, p=0.004) between treatments with Tukey post-hoc testing indicating significant differences between survival rates of G3 and G9 offspring compared to WT-derived offspring with the same father (Fig 3a). Offspring of G12 and the G8 F1 group both had lower survival than wild types with the same father, although these were not statistically significant. However, if individual G8 lines were compared, G8.4 had significantly lower survival than wild types (data not shown). The better survival in G8.5, 8.6 and 8.7 could be due to mosaicism in these females as they did not produce the expected number of transgenic offspring (see Table 3).This suggests that the degree of effect of a maternal G transgene on embryo early survival depends on transgene context.

Offspring of male frogs carrying G had much better survival to stage 14. Unpaired t-tests were used to compare survival rate for each male’s offspring to that of a wild type male crossed with the same female. No significant differences were found between pairs (Fig 3b), indicating no adverse effect on early embryogenesis when the father carries a G transgene. Early survival from each male’s offspring when crossed to the same wild type female (Fig 3c) also showed no significant variance between crosses (One-way ANOVA, F=1.05, p=0.44). In conclusion, paternal G did not affect survivability up to stage 14.

### Transgenic ratios are not skewed in G offspring

When the surviving embryos were 6 days old we were able to score them as transgenic or non-transgenic due to the presence or absence of red fluorescent protein RFP in the lenses of the eyes. χ-squared analysis was used to test for any deviation from expected Mendelian ratios, assuming a single transgene insertion site (i.e. founders are heterozygous at a single locus) (Table 3, significant p values indicate deviation). The survival rate for G9 offspring was too low to be meaningful for testing, but appeared Mendelian. G3 and G12 founder females, as well as the G8.4 female, showed the expected ratio (50:50) of transgenic offspring. G8.5, G8.6 and 8.7 generated a higher than expected number of wild types suggesting that these three females might be mosaic for the transgene. All female G8.x are derived from the G8 male, which showed typical Mendelian ratios of transgenic offspring (45% transgenic). None of the male frogs except G14 yielded non-Mendelian ratios of transgenic offspring (39% transgenic). The balance of transgenic and non-transgenic surviving embryos indicates that maternal, not zygotic, transgene status is responsible for the reduced survival of embryos from G mothers.

### Developmental progression up to 10 weeks is not affected by the presence of Gtransgene or by paren-t-of-origin

At two weeks, surviving offspring were transferred to a recirculating aquarium at 22°Cfor 8 weeks, to determine the effect, if any, of tadpole transgene presence, absence or source (i.e. maternal or paternal inheritance). Wild type tadpoles were derived from G3, G9, G12 females and G5 and G7 males; G female founder group were transgenic animals from G3, G9 and G12 mothers; G8.x founder group were transgenic animals from G8.4, 8.5, 8.6, 8.7 mothers, and male founders were transgenic animals from G5, G6, G7, G8 and G14 fathers. There was no difference in median developmental stage at 10 weeks of development: in each group, the median stage was 54 (Fig 3d). One-way ANOVA confirmed that there was no significant variation between groups. Hence, neither the origin or the presence/absence of a G transgene affected developmental stage when tadpoles were reared in normal aquarium temperatures (22°C).

**Table 3.** Transgene transmission tracking to offspring of female or male G heterozygote frogs. Deviations from expected Mendelian ratio of 50:50 generate a significant p value (bold) (Chi-squared test).

| Frog with transgene | Dead | % Wild type | % Transgenic | Chi squared | P-value |
| --- | --- | --- | --- | --- | --- |
| G3 female | 68 | 15 | 17 | 0.11 | 0.74 |
|  | 53 | 26 | 22 |  |  |
|  | 84 | 9 | 8 |  |  |
| G12 female | 86 | 6 | 7 | 0.03 | 0.85 |
|  | 91 | 6 | 3 |  |  |
|  | 89 | 5 | 6 |  |  |
| G8.4 female | 65 | 21 | 14 | 3.86 | 0.05 |
|  | 46 | 39 | 16 |  |  |
|  | 61 | 19 | 19 |  |  |
| G8.5 female | 2 | 74 | 23 | 88.6 | <0.0001 |
|  | 0 | 69 | 31 |  |  |
|  | 0 | 94 | 6 |  |  |
| G8.6 female | 16 | 65 | 19 | 42.12 | <0.0001 |
|  | 96 | 3 | 1 |  |  |
|  | 12 | 69 | 19 |  |  |
| G8.7 female | 18 | 58 | 24 | 6.91 | <0.01 |
|  | 45 | 35 | 19 |  |  |
|  | 30 | 39 | 30 |  |  |
| G5 male | 2 | 49 | 49 | 1.00 | 0.31 |
|  | 0 | 45 | 55 |  |  |
|  | 13 | 35 | 52 |  |  |
| G6 male | 16 | 38 | 46 | 0.11 | 0.74 |
|  | 18 | 42 | 40 |  |  |
|  | 4 | 47 | 49 |  |  |
| G7 Male | 11 | 50 | 39 | 1.94 | 0.16 |
|  | 6 | 62 | 32 |  |  |
|  | 9 | 43 | 48 |  |  |
| G8 Male | 18 | 44 | 38 | 1.60 | 0.21 |
|  | 15 | 48 | 37 |  |  |
|  | 6 | 50 | 44 |  |  |
| G14 male | 26 | 43 | 30 | 5.05 | 0.02 |
|  | 46 | 34 | 20 |  |  |
|  | 24 | 45 | 31 |  |  |

**Fig. 3.**
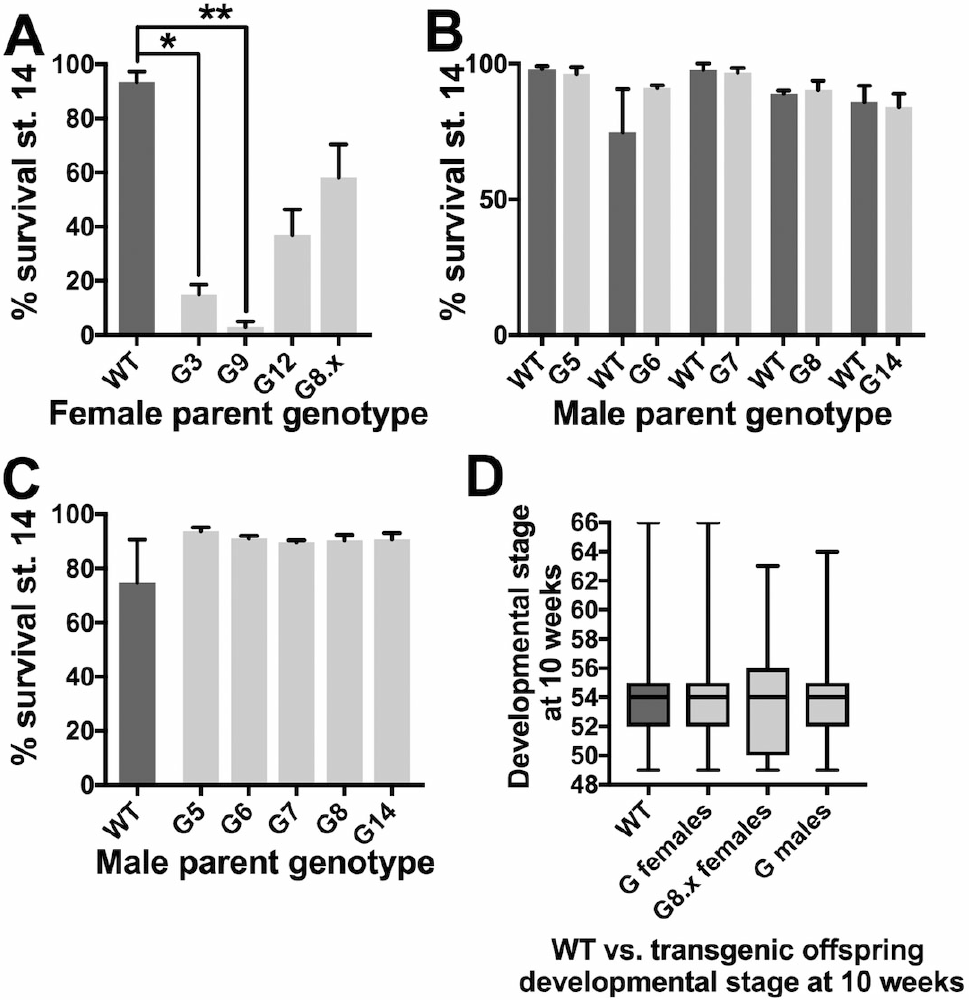
The low number of offspring from G females results from poor survival in very early embryonic development. A-C Column graphs showing the percentage of fertilized embryos that survive to stage 14 (the end of gastrulation). Columns represent the means of 3 replicate fertilisations of around 100 embryos each, and error bars are SEM. A) Offspring of females carrying a G transgene do not survive as well as those from WT female even when the father is the same. One-way ANOVA with Tukey post hoc pairwise testing showed G3 (p<0.05, *) and G9 (p<0.01, **) have significantly lower survival compared to wild types. B) No differences were observed between early stage survivability in offspring of five different male G founder frogs,when compared to the offspring of WT males with the same mother. C) Similarly, when all five G males were crosses to the same wild type female, early development was unaffected. D) Box plot of developmental stage at 10 weeks post fertilization for offspring grouped by genotype. Median developmental stage is depicted by the line, with boxes representing first and third quartile, and whiskers showing the minimum and maximum stage, up to stage 66 which is a fully metamorphosed juvenile frog. Median stage at 10 weeks was unaffected by the G transgene regardless of parental or individual genotype. Genotype sample sizes were: WT n=145, G transgenics derived from carrier females n=132, G transgenics derived from G8.x females n=47 and G transgenics derived from G males n=99.

### Ovarian recovery is not affected by the *grem1* transgene

Induction of *Xenopus* female adults with human chorionic gonadotrophin (HCG) normally results in the maturation and laying of all stage VI oocytes. All female *grem1* (G) frogs in the study were sacrificed at 6 weeks after induction and the ovaries extracted. This time is expected to allow for full regeneration, replenishing stage VI oocytes (13).

A single lobe from each ovary was isolated for each female and treated with collagenase to separate the oocytes. Oocytes were than staged and counted according to Dumont as stage II to VI, or atresic (dying). Some stage I oocytes were also observed but as many were lost on collagenase treatment, they could not be reliably counted (13).

Interestingly, frogs were weighed before sacrifice and found to have lost around 25% of their bodyweight compared to pre-induction. Thus, even after 6 weeks, the ovary is not fully recovered as suggested by Dumont. This was also reflected in the percentage of stage VI oocytes, shown in Fig 4a. While a female wild type that had not been induced for 1 year had 58% oocytes at stage VI, induced wild types had 19 and 20% respectively, with G founders ranging from 16 to 36% and G8 F1 females 29-32%. Data were grouped into G founders (F0), G8 F1 and WT, and analysed by one way ANOVA, no significant variance was detected (F=1.42, p=0.27). However, if the wild type female that had not been induced 6 weeks previously was included, a significant difference could be detected between this animal and the other study groups (Fig 4a). This suggested that ovarian regeneration was incomplete, but similar, in transgenic and non-transgenic females, provided the regeneration time was consistent. Histological and ultrastructural analysis of wild type vs. transgenic ovary samples showed no obvious phenotypic differences in the ovary (data not shown).

**Fig. 4.**
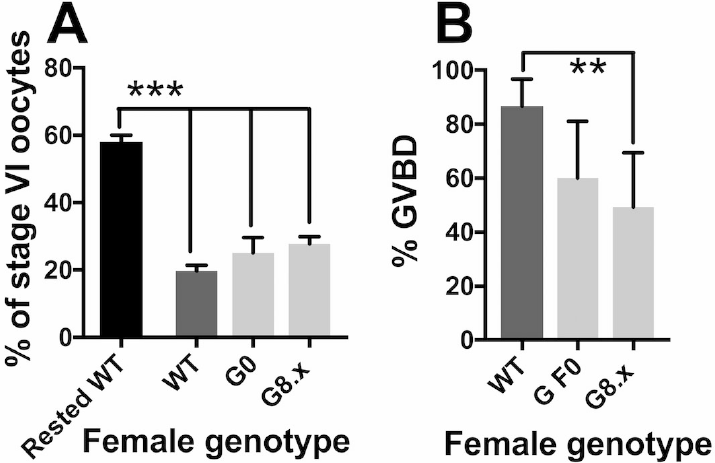
Ovarian recovery and maturation of stage VI oocytes is normal in G female frogs. Column graphs showing mean and error bars showing SEM. Grouped transgenic sample sets as for previous figures. A) The %age of stage VI oocytes in a randomly chosen left and right ovary lobe from WT (n=2), G founders (n=3) or G8 F1 (n=4) generation female frogs, representing the amount of ovarian recovery. The presence of the maternal transgene did not significantly the %age of stage VI oocytes when compared with wild types that had been induced at the same time (6 weeks prior). However, none of the ovaries in the study had fully regenerated 6 weeks after induction. “Rested WT” shows comparative data from a WT female last induced at > 1 year prior to sacrifice, and all 3 groups had significantly fewer stage VI oocytes than this female (1 way ANOVA with post hoc Tukey test, p<0.0010***). B) %age of stage VI oocytes responding appropriately to progesterone treatment, scored by the presence of the germinal vesicle as a white spot at the animal pole of the oocyte, and indicating successful maturation in vitro. Fisher’s exact test showed significantly fewer G8.x oocytes responded to progesterone compared to WT (p<0.01 **).

### Response of stage VI oocytes to progesterone is unaffected by the presence of a G transgene

Stage VI oocytes are arrested in G2/M meiotic prophase I, and cannot be fertilised. Exposure to progesterone initiates progression to metaphase II and germinal vesicle breakdown (the breakdown of the egg nuclear membrane, termed GVBD). Oocytes that have undergone GVBD have a white spot at the animal pole, resulting from displacement of pigment from the egg cortex by the nucleus. Some of the stage VI oocytes from G females appeared less responsive to progesterone (Fig 4b), and Fisher’s exact test showed G8.x oocytes to be significantly less likely to mature on progesterone exposure (p=0.0035). Grouping all G female oocytes and comparing to WT also revealed a significant reduction in numbers of matured oocytes (Fisher’s exact test, p=0.0137).

### Q-RT-PCR reveals a reduced level of *grem1* transcripts in the ovary of G founders and G8 F1 compared to that of non-transgenics

Initially, we thought that *grem1* might be transiently activated from the *hsp70:grem1*, γ-*crystallin:RFP* (G) transgene during oogenesis, leading to poor development of offspring from G transgene carrying mothers. Previous work has shown that endogenous hsp70 is heat inducible, but normally silent in stage VI oocytes and constitutively active in younger, vitellogenic oocytes (20). This suggests that the heat shock factors that bind to and activate the endogenous *hsp70* locus could also be driving expression from the transgene. To determine if *grem1* was being expressed from the *hsp70* transgenic promoter, we used qRT-PCR of total RNA extracted from wild type and transgenic oocytes and ovaries, harvested 6 weeks after initial induction of egg laying. As primers cannot not distinguish between endogenous *grem1* and transgene *grem1* transcripts, this assay measures both together. Unexpectedly, we saw a consistently lower *grem1* expression in stage I to VI oocytes from G females when compared to their wild type counterparts (Fig 5a). Significant reduction in *grem1* transcripts was seen at stage V and VI (Bonferroni multiple comparisons test for WT and G at each oocyte stage). To further investigate this surprising result, we analysed *grem1* expression in both ovaries of every frog in the study. Collectively, both the G0 founders and the G8.x F1 female ovaries expressed significantly lower *grem1* than wild types (Fig 5c, 1 way ANOVA, with Tukey post-hoc testing p<0.0001). This was due to more than 2-fold down regulation of *grem1* in all the lines (Fig 5b), with G9 the most affected at almost 5-fold down regulated. Interestingly, G9 showed the poorest survival rate (refer to Fig 2a), with just 3% of embryos surviving past gastrulation. Surprisingly then, the presence of the transgene in the mother results in a decrease in grem1 transcripts. This suggests that the endogenous *grem1* locus is less active in the presence of the G transgene. To see if endogenous hsp70 was influenced by the transgene, we also measured hsp70 transcripts but found no difference in expression between wild type and transgenic ovaries (Fig 5d, 1 way ANOVA, Tukey post-hoc testing). Since activation of the transgene should also result in activation of the endogenous locus, increasing *hsp70* transcripts, this suggests that the transgene is not active, or leaky, in the ovary, and that there is no significant effect of transgene presence on the endogenous *hsp70* locus, at least in this tissue.

To look at another locus for comparison, we measured *bmp15* transcripts, as BMP15 levels in follicular fluid are a well-documented indicator of oocyte quality and successful early development in humans. *bmp15* expression varied between the three groups (Fig 5e, 1 way ANOVA F=6.33, p=0.003) and was significantly higher in G8.x compared to either G0 founders or WT ovaries, (Tukey post hoc-testing, p=0.009 and 0.010 respectively). However we did not observe a strong correlation between *bmp15* and *grem1* transcript levels across all samples, indicating that the reduced survival ofG female offspring is not due to altered *bmp15* expression.

**Fig. 5.**
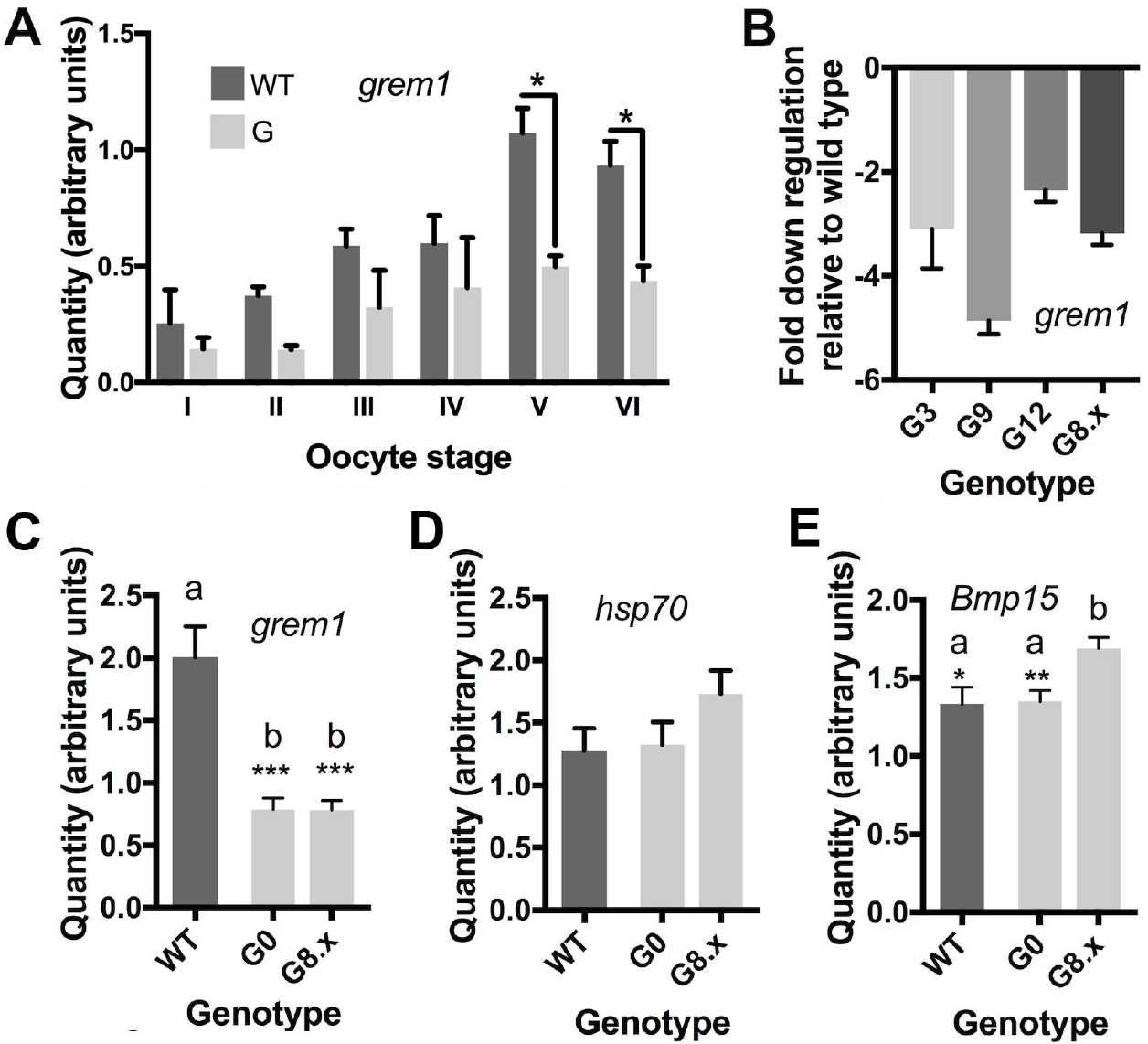
Q-RT-PCR revealed a decrease in whole ovary grem1 that correlates with poor survivability of the offspring of G mothers. A-E) Column graphs of transcript levels measured by q-RT-PCR for oocytes (A) or a randomly chosen left and right ovary lobe (B-E) from each animal. Samples were tested in triplicate and normalised to the geometric mean of two housekeepers, Gapdh and ef-1α, except in A where normalisation was to Gapdh. In B, columns show means of relative quantity grem1 transcripts in G ovaries relative to wild type. In C-E, columns depict means of technical and biological replicates for each group of genotypes and with error bars are SEM. Quantities are expressed as arbitrary values calculated from standard curves. A) *grem1* is expressed at lower levels in oocytes from G females at all stages of oocyte development, this is significant for stage V and VI oocytes. Data is the mean of triplicate data from two G females, G8.4 and G3, and two wild type (WT) females. B) *grem1* transcripts are at least 2 fold lower in G ovaries than in wild type, with G9 (which had the poorest survival) having nearly 5-fold reduction in *grem1* transcripts. C) *grem1* is expressed at significantly lower levels than WT in either G founders or G8.x ovaries. C) *hsp70* transcripts are not significantly different between WT and G ovaries. D) bmp15 transcript levels were significantly higher in the G8 F1 ovaries, than either WT or G founders. * p<0.05, **p <0.01, ***p<0.001

## Discussion

### *grem1* may be a pan-vertebrate marker of oocyte competence

In humans, the success of assisted fertility depends on being able to predict oocyte viability (Reviewed by (21)). The competence of the oocyte to mature, fertilise, develop and implant into the uterine wall will determine the rate of successful pregnancy. It is highly desirable to be able to identify the most competent oocytes to avoid having to transfer multiple embryos and risk multiple pregnancies. Oocyte competence is known to be closely associated with normal follicular development, with intrinsic reciprocal signals between the oocyte and its surrounding follicle cells playing as key role, as well as extrinsic, circulating hormonal factors. Here, we have shown a correlation between maternal *grem1* and early embryonic survival in the highly fecund frog *Xenopus laevis*. Although we cannot rule out an effect of the presence of the transgene in mothers on extrinsic hormone regulation, our results favour an intrinsic effect on ovarian follicle cells. The presence of the G transgene in follicle cells results in significantly lower levels of *grem1* transcripts in the ovary, suggesting that the transgene somehow results in repression of the endogenous *grem1* gene. Lower levels of *grem1* transcripts would be expected to reduce available *grem1* protein, potentially resulting in de-repression of maternal BMP4/7 signalling. While the exact mechanism remains unknown, candidate gene approaches have shown that grem1 transcript abundance is also a predictor of human oocyte competence (16, 17). In an unbiased microarray study, Assidi et al also identified *grem1* as one of a handful of markers associated with oocyte competence in a bovine model (22). Taken together with our results in the frog model, *grem1* levels seem to be a marker of oocyte competence in highly fecund non-mammalian as well as mammalian vertebrates. An exception to this is the mouse, where conditional, ovar-specific *grem1* null mothers showed normal fertility (23). A possible explanation for this is that *grem2*, which is also expressed in the follicle cells, may compensate for loss of *grem1* in mouse ovaries. *Grem2*, which is not found in *Xenopus*, seems have a conserved role in regulation of primordial follicle transition in some mammals (24, 25). However, in mice, grem1 has been shown to play this role (23). Some flexibility in the roles of the two orthologues therefore exists in mammals.

### Maternal presence of the G transgene in *Xenopus* results in poor survival in early embryogenesis

The most likely explanation for the reduced *grem1* in ovaries of G female frogs is that the presence of the transgene is somehow supressing the transcription of the endogenous *grem1* locus. This effect was seen to varying degrees in different lines, suggesting that integration position has some effect on the amount of repression. *grem1* transcripts were consistently reduced, relative to wild type controls, at all stages of oocyte development (data not shown) and in all 4 lines tested. Transgene silencing has been studied extensively in plants, where its discovery helped elucidate mechanisms of epigenetic silencing such as RNAi. In plants, two types of silencing are known (reviewed in (26)). Position effects, where the transgene is negatively regulated by flanking DNA or location in heterochromatic regions, seem unlikely to be the cause of the silencing we observe here, since each line would be expected to have the transgene integrated in a different random location. Further more,the *hsp70:grem1* transgene is linked to γ*Crys:RFP*, allowing identification of transgenic animals without genotyping, and clearly this is not silenced. The second mechanism is termed homology-dependent gene silencing (HDGS) which may result from multiple transgenes integrating at one site in random orientation, which may result in inverted repeats that can interact with, and silence, homologous sequences in the genome. HDGS seems the more likely explanation, since it could explain why *grem1* transcripts, but not *hsp70* transcripts, are significantly reduced. Similar transgene silencing observed in mammals has been shown to arise from inverted repeats of a transgene (27) although the mechanism was not able to be fully characterised.

## Conclusions

Female frogs carrying a heat-inducible transgene driving conditional expression of *grem1* produce few viable offspring. Here we have shown that this is not due to reduced fecundity or fertilization rate, but that the offspring of these females die in early embryogenesis regardless of whether of not they themselves carry the transgene. This maternal effect lethality is not caused by transgene leakage, which would elevate *grem1* expression, as we observe reduced levels of *grem1* transcripts in the *Xenopus* ovary of transgenic mothers. Therefore, it is likely *grem1* levels are associated with chance of survival in the early stages of embryogenesis in frogs as they have been found to in mammals (16,17, 21). Furthermore, this indicates that transgene suppression of an endogenous locus may occur in the *Xenopus* female germline, an unexpected finding. The mechanism by which this occurs is presently unknown, but could be a case of homology-dependent gene silencing.

## ACKNOWLEDGEMENTS

CWB conceived the study, designed the experiments, analysed data and wrote the manuscript. LT and CWB performed all *Xenopus* breeding and experiments and CB and JW performed all qRT-PCR analysis. All authors read and approved the final manuscript. We are grateful to Matthew Downes for his expertise in electron microscopy studies even though this data did not make the final cut. Mark Lokman kindly read and commented on a draft and provided a second opinion on statistics. Jolyn Chia, Nikita Woodhead and Nat Lim take excellent care of our frog colony, and Valerie Gavois arranged the internships for CB and LT who are students at Ecole Nationale Vétérinaire de Toulouse, France.

